# Valid post-clustering differential analysis for single-cell RNA-Seq

**DOI:** 10.1101/463265

**Authors:** Jesse M. Zhang, Govinda M. Kamath, David N. Tse

## Abstract

Single-cell computational pipelines involve two critical steps: organizing cells (clustering) and identifying the markers driving this organization (differential expression analysis). State-of-the-art pipelines perform differential analysis after clustering on the same dataset. We observe that because clustering *forces* separation, reusing the same dataset generates artificially low *p*-values and hence false discoveries. We introduce a valid post-clustering differential analysis framework which corrects for this problem. We provide software at https://github.com/jessemzhang/tn_test.

## Introduction

Modern advances in single-cell technologies can cheaply generate genomic profiles of millions of individual cells [1, 2]. Depending on the type of assay, these profiles can describe cell features such as RNA expression, transcript compatability counts [3], epigenetic features [4], or nuclear RNA expression [5]. Because the cell types of individual cells often cannot be known prior to the computational step, a key step in single-cell computational pipelines [6, 7, 8, 9, 10] is clustering: organizing individual cells into biologically meaningful populations. Furthermore, computational pipelines use differential expression analysis to identify the key features that distinguish a population from other populations: for example a gene based on its relative expression level.

Many single-cell RNA-seq discoveries are justified using very small *p*-values [9, 11]. The central observation underlying this paper is: these *p*-values are often *spuriously* small. Existing workflows perform clustering and differential expression on the same dataset, and clustering *forces* separation regardless of the underlying truth, rendering the *p*-values invalid. This is an instance of a broader phenomenon, colloquially known as “data snooping”, which causes false discoveries to be made across many scientific domains [12]. While several differential expression methods exist [9, 13, 14, 11, 15, 16], none of these tests correct for the data snooping problem as they were not designed to account for the clustering process. As a motivating example, we consider the classic Student’s *t*-test introduced in 1908 [17], which was devised for controlled experiments where the hypothesis to be tested was defined before the experiments were carried out. For example, to test the efficacy of a drug, the researcher would randomly assign individuals to case and control groups, administer the placebo or the drug, and take a set of measurements. Because the populations were clearly defined a priori, a *t*-test would yield valid *p*-values. In other words, under the null hypothesis where no effect exists, the *p*-value should be uniformly distributed between 0 and 1. For single-cell analysis, however, the populations are often obtained, via clustering, *after* the measurements are taken, and therefore we can expect the *t*-test to return significant *p*-values even if the null hypothesis was true. The clustering introduces a **selection bias** [18, 19] that would result in several false discoveries if uncorrected.

## Results

In this work, we introduce a method for correcting the selection bias induced by clustering. To gain intuition for the method, consider a single-gene example where a sample is assigned to a cluster based on the expression level of the gene relative to some threshold. Fig. 1a shows how this expression level is deemed significantly different between two clusters even though all samples came from the same normal distribution. We attempt to close the gap between the blue and green curves in the rightmost plot by introducing the truncated normal (TN) test. The TN test (Fig. 1b) is an approximate test based on the truncated normal distribution that corrects for a significant portion of the selection bias. As we go from 1 gene to multiple, the decision boundary generalizes from a threshold to a high-dimensional hyperplane. We condition on the clustering event using the hyperplane that separates the clusters, and Table 1 shows that this linear separability assumption is valid for a diverse set of published single-cell datasets. By incorporating the hyperplane into our null model, we can obtain a uniformly distributed *p*-value even in the presence of clustering. To our knowledge, the TN test is the first test to correct for clustering bias while addressing the differential expression question: *is this feature significantly different between the two clusters?* We then proceed to provide a data-splitting based framework (Fig. 1c) that allows us to generate valid differential expression *p*-values for clusters obtained from any clustering algorithm. Using both synthetic and real datasets, we argue that in the existing framework, not all reported markers can be trusted for a given set of clusters. This point implies that

1. large correction factors for multiple markers can indicate overclustering;
2. plotting expression heatmaps where rows and columns are arranged by cluster identity can convey misleading information (e.g. Fig. S6 in [20] and Fig. 6b in [1]).

**Figure 1:**
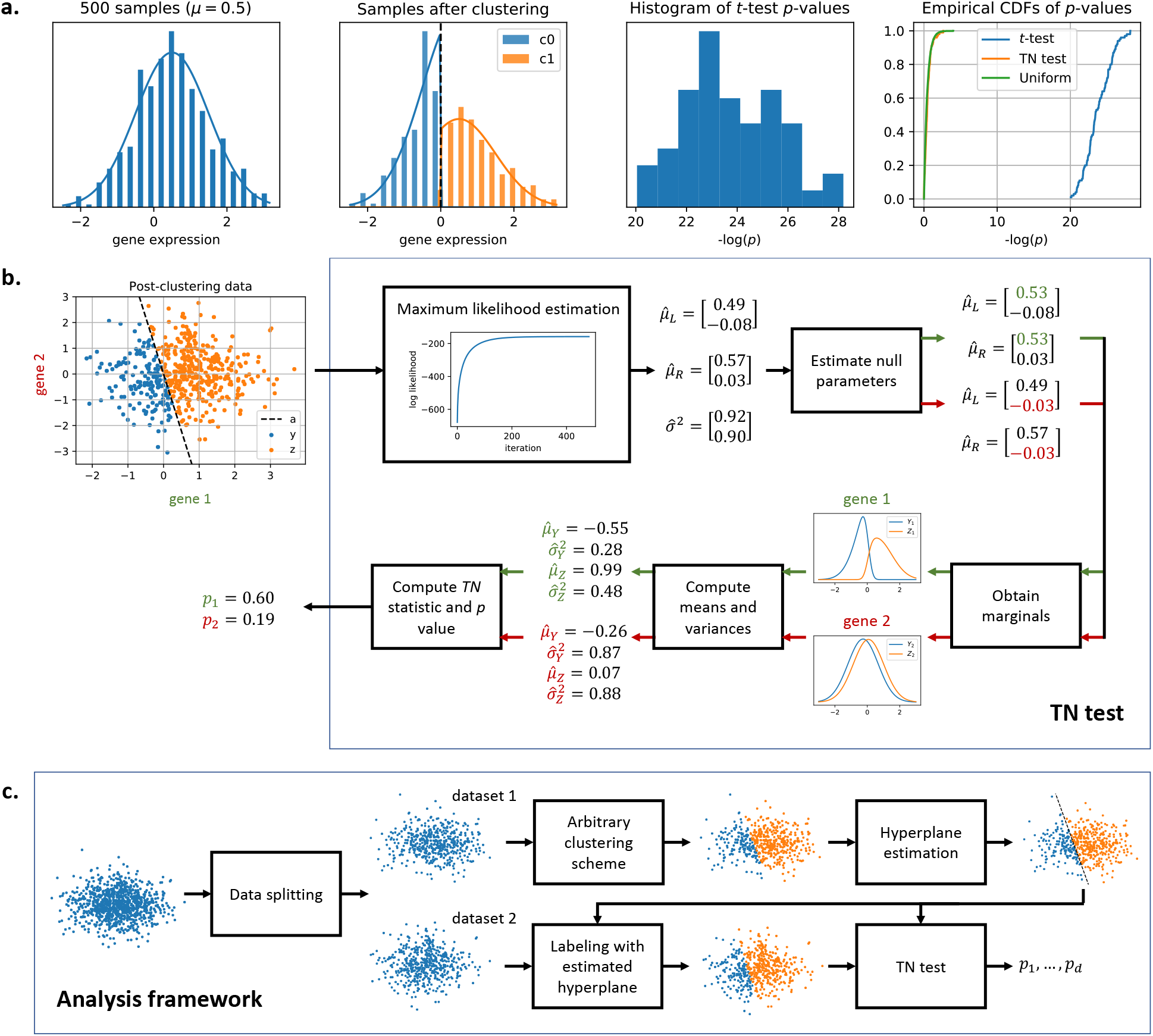
Overview of the truncated normal (TN) test. **a.** Although the samples are drawn from the same normal distribution, a simple clustering approach will always generate two clusters that seem significantly different under the *t*-test. The TN test statistic is based on two truncated normal distributions where the truncation is at the clustering threshold. This allows the test to account for and correct the selection bias due to clustering, closing the gap between the blue and green cumulative distribution functions (CDFs) in the rightmost plot. **b.** *TN test for differential expression. μ_L_* and *μ_R_* are the means of the untruncated normal distributions that generated *Y* and *Z*. The covariance matrix is assumed to be diagonal and equal across the two untruncated distributions, and *σ*^2^ represents the diagonal entries along the matrix. The threshold in the singlegene case generalizes to a separating hyperplane *a* in the multi-gene case. The TN test assumes the hyperplane *a* is given. **c.** *Analysis framework*. The samples are split into two datasets. The analyst’s chosen clustering algorithm is performed on dataset 1, and a hyperplane is fitted to the cluster labels. The hyperplane, which is independent from dataset 2, is then used to assign labels to dataset 2. Differential expression analysis is performed on dataset 2 using the TN test with this separating hyperplane.

**Table 1:** Training and validation accuracies obtained after fitting one-v-rest logistic regression models on various published single-cell RNA-Seq datasets. The predicted labels were provided by the authors. For each dataset, the samples were divided into five folds. Four folds (80% of the samples) were used for training, and the remaining fold (20% of the samples) was used for validation. The reported accuracies are averaged across all five folds.

| dataset | # of samples | # of classes | train accuracy | valid accuracy |
| --- | --- | --- | --- | --- |
| 10x [2] | 39999 | 9 | 0.96 +/- 0.00 | 0.95 +/- 0.00 |
| Biase [29] | 49 | 3 | 1.00 +/- 0.00 | 0.96 +/- 0.04 |
| Birey [30] | 11835 | 14 | 1.00 +/- 0.00 | 0.92 +/- 0.00 |
| Buettner [31] | 182 | 3 | 1.00 +/- 0.00 | 0.65 +/- 0.02 |
| Deng [32] | 264 | 9 | 1.00 +/- 0.00 | 0.97 +/- 0.01 |
| DropSeq [1] | 44808 | 39 | 0.99 +/- 0.00 | 0.96 +/- 0.00 |
| Joost [33] | 1422 | 13 | 1.00 +/- 0.00 | 0.81 +/- 0.01 |
| Kolodziejczyk [27] | 704 | 3 | 1.00 +/- 0.00 | 1.00 +/- 0.00 |
| Patel [34] | 430 | 5 | 1.00 +/- 0.00 | 0.88 +/- 0.02 |
| Pollen [35] | 249 | 11 | 1.00 +/- 0.00 | 0.98 +/- 0.01 |
| Ting [36] | 149 | 5 | 1.00 +/- 0.00 | 0.96 +/- 0.01 |
| Treutlein [37] | 77 | 4 | 1.00 +/- 0.00 | 0.84 +/- 0.02 |
| Usoskin [38] | 696 | 5 | 1.00 +/- 0.00 | 0.91 +/- 0.01 |
| Yan [39] | 118 | 6 | 1.00 +/- 0.00 | 1.00 +/- 0.00 |
| Zeisel [20] | 3005 | 9 | 1.00 +/- 0.00 | 0.96 +/- 0.01 |

We validate the TN test framework on synthetic datasets where the ground truth is fixed and known (see STAR Methods). We first demonstrate the performance of the proposed framework on real datasets before expanding on method details.

### Performance on single-cell RNA-Seq data

We consider the peripheral blood mononuclear cell (PBMC) dataset of 2700 cells generated using recent techniques developed by 10x Genomics [2], and this dataset was also used in a tutorial for the Seurat single-cell package [6]. The Seurat pipeline yielded 9 clusters output after preprocessing the dataset and running a graph-based clustering algorithm [21, 22, 23]. For each of 7 approaches offered by Seurat (see STAR Methods for more details), we perform differential expression analysis on clusters 0 and 1 (Fig. 2a), which are T-cell subtypes according to the Seurat tutorial [6], and we compare the obtained *p*-values to TN test *p*-values. Fig. 2a shows that while the TN test agrees with the differential expression tests on several genes (e.g. *S100A4*), it also disagrees on some other genes. The two genes with the most heavily corrected *p*-value were *B2M* and *HLA-A*. While several of the Seurat-provided tests would detect a significant change (e.g. the popular Wilcoxon test reported *p* = 8.5 × 10^−30^ for *B2M* and *p* = 3.8 × 10^−17^ for *HLA-A*), the TN-test accounts for the fact that this difference in expression may be driven by the clustering approach (*p* = 3.7 × 10^−13^ for *B2M* and *p* = 7.2 × 10^−8^ for *HLA-A*). Reads from *HLA-A* are known to generate false positives due to alignment issues [24]. Because the amount of bias correction is different for each gene, the TN test orders markers differently than clustering-agnostic methods. Notably, the TN test identifies two gene markers missed by the other tests: *ANXA1*, which is associated with T-cells [25], and *S100A6*, which is associated with whole blood [26]. Comparisons are also performed for clusters 1 versus 3 and 2 versus 5, and the results are reported in Supplementary Fig. 2.

**Figure 2:**
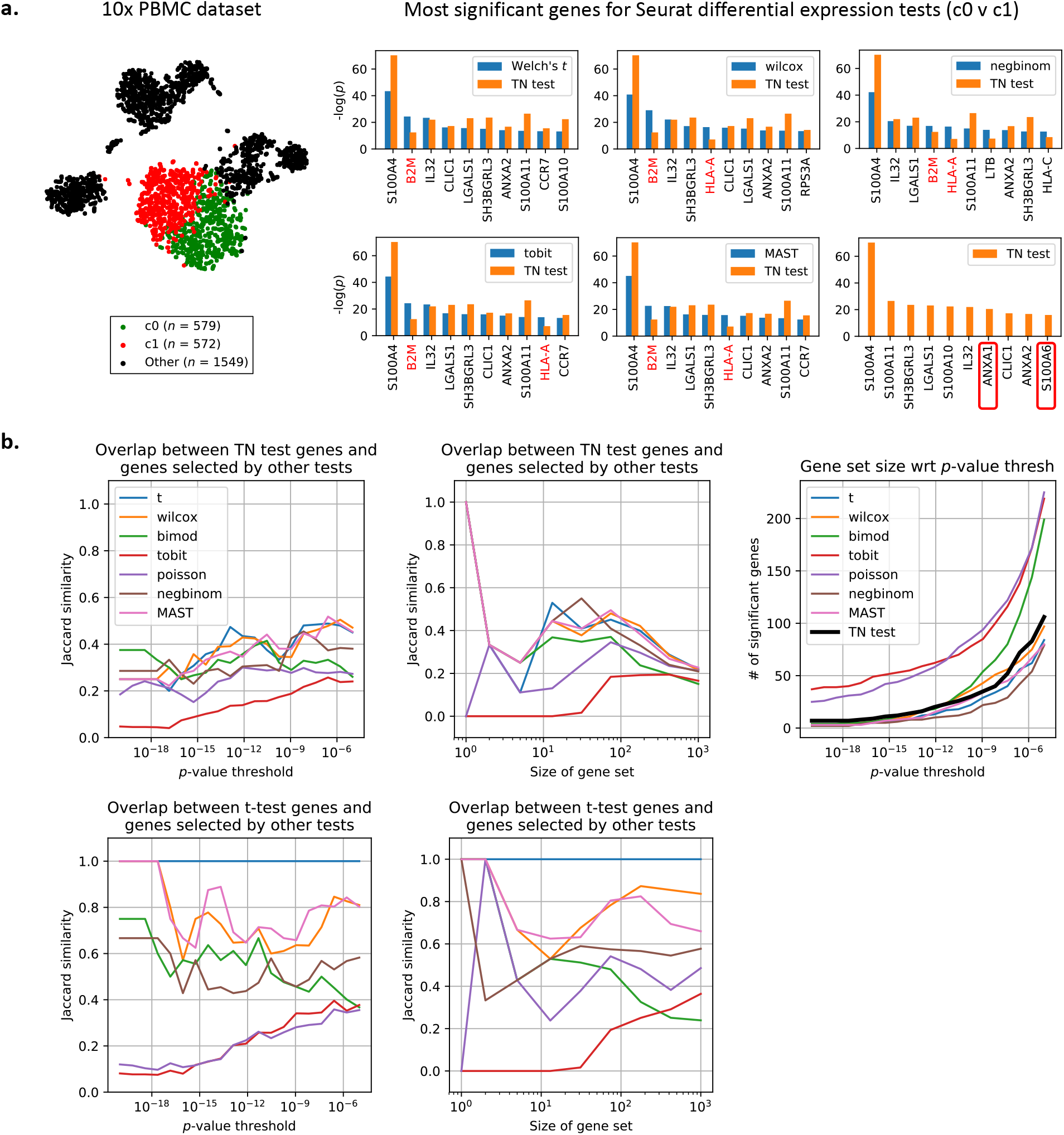
Comparison of TN test to 7 other tests on peripheral blood mononuclear cell (PBMC) dataset. **a.** t-SNE plot of the 2700-PBMC dataset with two clusters found using Seurat [6]. Following the analysis pipeline in Fig. 1, the Seurat clustering approach is used to recover 2 clusters on dataset 1, and seven differential expression methods provided by Seurat (see STAR Methods for details) are also run on dataset 1. The TN test ranks genes differently than standard differential expression methods, indicating that artifacts of post-selective inference are consequential in picking relevant genes. The TN test also identifies two markers (boxed in red) missed by the other tests. **b.** The TN test and *t*-test are compared to the 7 other tests using a variety of metrics. For more details on Jaccard similarity, see STAR Methods.

We compare the genes selected by the TN test to those selected by the 7 other approaches using a variety of metrics. Fig. 2b shows that while the TN test agrees with a fraction of selected genes for all *p*-value thresholds tested, in general the TN test does not agree with the tests as much as the tests agree with each other. This again emphasizes how none of the 7 tests account for the clustering selection bias, and therefore they should make similar mistakes (i.e. fail to correct some subset of genes). Fig. 2b (right-most panel) also shows how the TN test returns a comparable number of genes to other state-of-the-art tests. We can visualize the overlap between the top 100 genes chosen by the TN test compared to those chosen by the Welch’s *t*-test using a permutation matrix (Supplementary Fig. 3). If the same set of genes were selected in both sets, then the permutation matrix would be diagonal. Fig. 2 indicates that the artifacts of post-selection inference are fairly consequential even in real datasets.

We further explore how the TN test can be used to both validate and contest reported subtypes. For a dataset of 3005 mouse brain cells [20], the authors reported 16 subtypes of interneurons using 26 gene markers. Fig. 3a and Supplementary Fig. 4 shows that Int11, the only subtype that was experimentally validated using immunohistochemistry, received relatively small amounts of correction. Int1, Int12, and Int16, however, may need further inspection.

**Figure 3:**
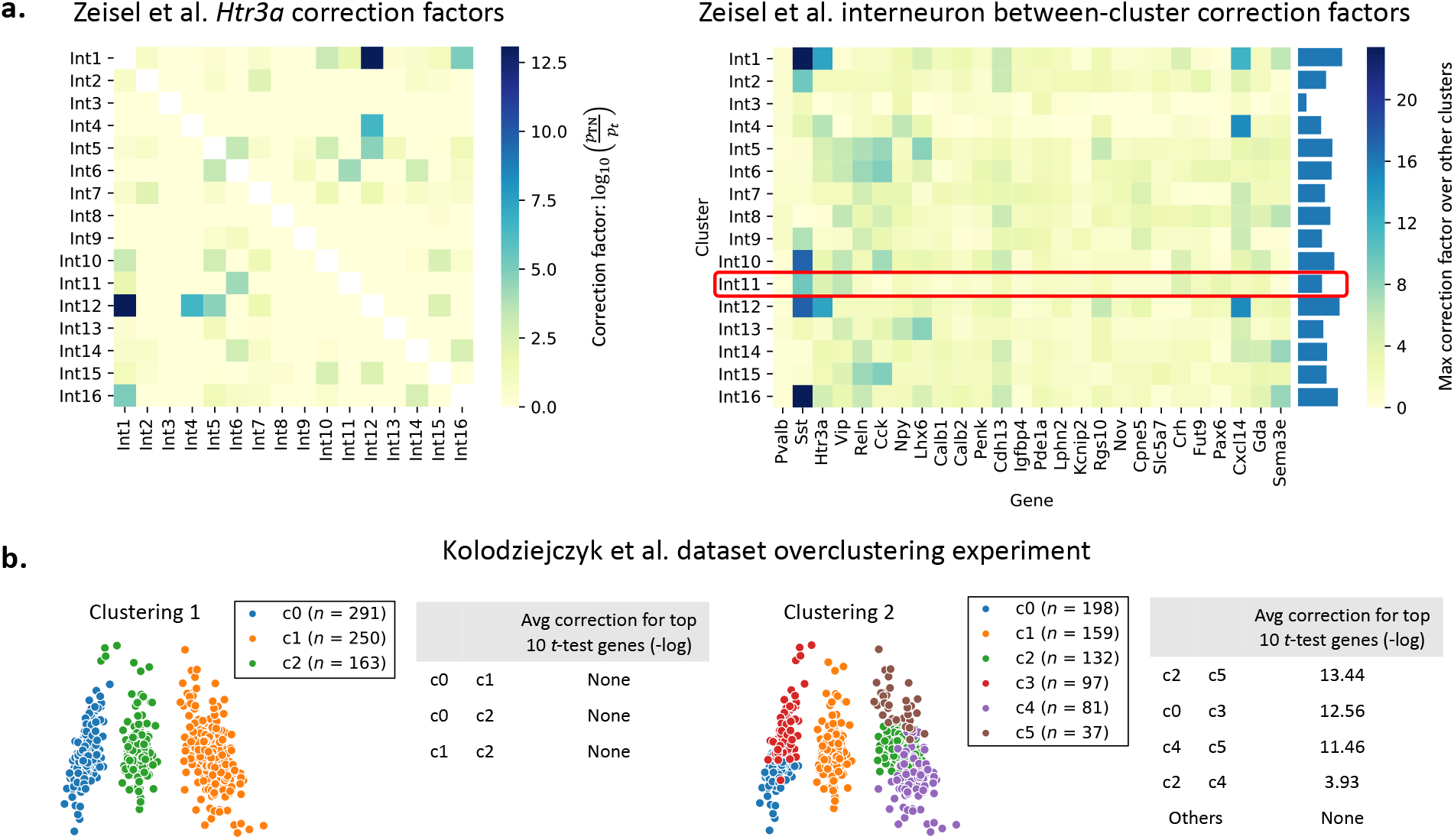
TN test on mouse brain cell and mESC datasets. **a.** The 16 interneuron subclasses reported for the mouse brain cell dataset [20] are re-compared using each of the 26 genes discussed by the authors. The left plot shows an example correction factor heatmap for one of the 26 genes; the rest are reported in Supplementary Fig. 4. The right plot shows the max correction factor for each cluster and gene across all other clusters. Cluster Int11 was the only cluster verified to have biological significance by the authors in [20]. **b.** The Seurat pipeline is run with two different clustering parameters for the mESC dataset [27]. “None” indicates that TN test *p*-values were on average at least as significant as *t*-test ones. Pairs of clusters which look separated (Clustering 1, left pane) undergo no significant correction while pairs of clusters that do not look well-separated (e.g. c2 and c5 in Clustering 2, right pane) undergo high correction.

We also demonstrate how the TN test can be used to gauge overclustering. We run the Seurat clustering pipeline on a dataset of 704 mouse embryonic stem cells (mESCs) [27] using two different clustering parameters, resulting in the two clustering results shown in Fig. 3b. For each pair of clusters, we look at the top 10 most significant genes chosen by the *t*-test. We correct these *p*-values using the TN test and observe the geometric average of the ratio of TN test *p*-values to *t*-test *p*-values. We see that for valid clusters (Clustering 1), the *p*-value obtained using the TN test is often even smaller, offering no correction. When clusters are not valid (Clustering 2), however, we observe a significant amount of correction.

### Framework details

Next we describe the technical details of the method we develop in this manuscript.

#### Clustering model

To motivate our approach, we consider the simplest model of clustering: samples are drawn from one of two clusters, and the clusters can be separated using a linear separator. For the rest of this section, we assume that the hyperplane *a* is given and independent from the data we are using for differential expression analysis. For example, we can assume that in a dataset of *n* independent and identically distributed samples and *d* genes, we had set aside *n*_1_ samples to generate the two clusters and identify *a*, thus allowing us to classify future samples without having to rerun our clustering algorithm. We run differential expression analysis using the remaining *n*_2_ = *n* − *n*_1_ samples while *conditioning* on the selection event. More specifically, our test accounts for the fact that a particular a was chosen to govern clustering. We will later demonstrate empirically that the resulting test we develop suffers from significantly less selection bias.

For pedagogical simplicity, we start by assuming that our samples are 1-dimensional (*d* = 1) and our clustering algorithm divides our samples into two clusters based on the sign of the (mean-centered) observed expressions. Let *Y* represent the negative samples and *Z* represent the positive samples. We assume that our samples come from normal distributions with known variance 1 prior to clustering, and we condition on our clustering event by introducing truncations into our model. Therefore *Y* and *Z* have *truncated* normal distributions due to clustering:

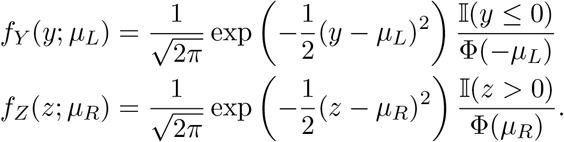

Here, the 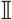 terms are indicator functions denoting how truncation is performed, and the Φ terms are normalization factors to ensure that *f_Y_* and *f_Z_* integrate to 1. Φ represents the CDF of a standard normal random variable. *μ_L_* and *μ_R_* denote the means of the untruncated versions of the distributions. We want to test if the gene is differentially expressed between two populations *Y* and *Z*, i.e. if *μ_L_* = *μ_R_*.

#### Derivation of the test statistic

The joint distribution of our *n* samples of *Y* with our *m* samples of *Z* can be expressed in exponential family form as

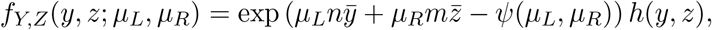

where *ȳ* and 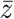 represent the sample means of *Y* and *Z*, respectively. *ψ* is the cumulant generating function, and *h* is the carrying density. Please see STAR Methods for more details. To test for differential expression, we want to test if *μ_L_* = *μ_R_*, which is equivalent to testing if *μ_R_ − μ_L_* = 0. With a slight reparametrization, we let *θ* = (*μ_R_ − μ_L_*)/2 and *μ* = (*μ_R_ + μ_L_*)/2, resulting in the expression:

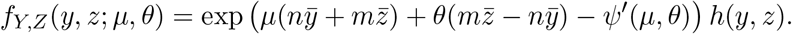

We can test design tests for *θ* = *θ*_0_ using its sufficient statistic, 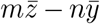 [28]. From the Central Limit Theorem (CLT), we see that the test statistic

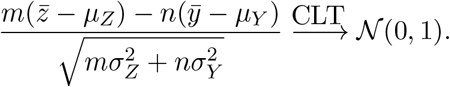

Intuitively, this test statistic compares 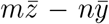, the gap between the observed means, to *mμ_Z_ − nμ_Y_*, the gap between the expected means. For differential expression, we set *θ*_0_ = 0. Because 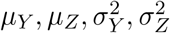 under the null *μ_L_* = *μ_R_* are unknown, we estimate them from the data by first estimating *μ_L_* and *μ_R_* via maximum likelihood. Although the estimators for *μ_L_* and *μ_R_* have no closed-form solutions due to the Φ terms, the joint distribution can be represented in exponential family form. Therefore the likelihood function is concave with respect to *μ_L_* and *μ_R_*, and we can obtain estimates 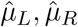 via gradient ascent. We then set 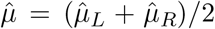. This procedure is summarized in Algorithm 1. We note that approximation errors are accumulated from the CLT approximation and errors in the maximum likelihood estimation process, and therefore the limiting distribution of the test statistic should have wider tails. Despite this, we show later that this procedure corrects for a large amount of the selection bias.

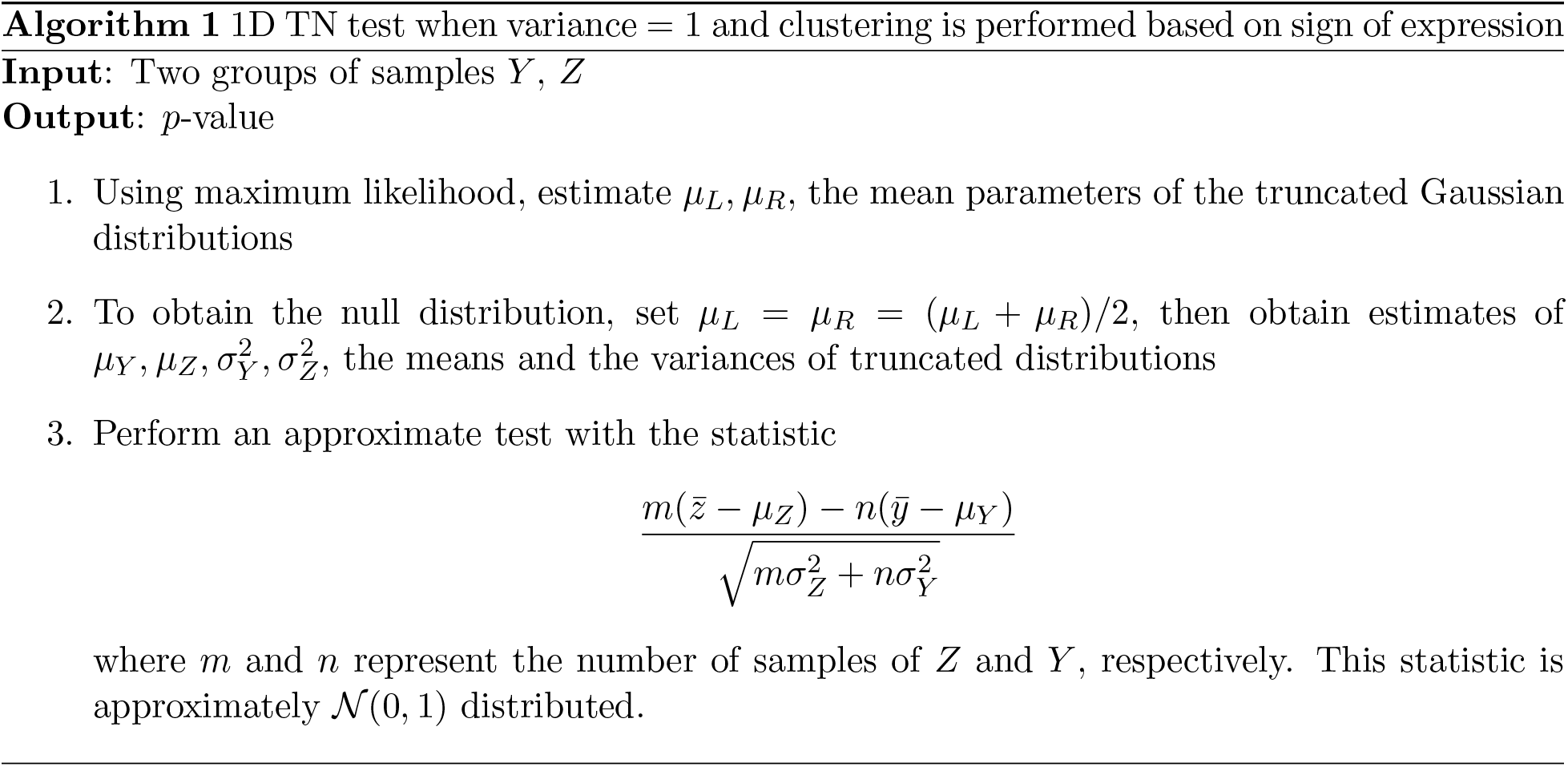

#### TN test for *d* dimensions and unknown variance

In this section, we generalize our 1-dimensional result to *d* dimensions and non-unit variance. Our samples now come from the multivariate truncated normal distributions

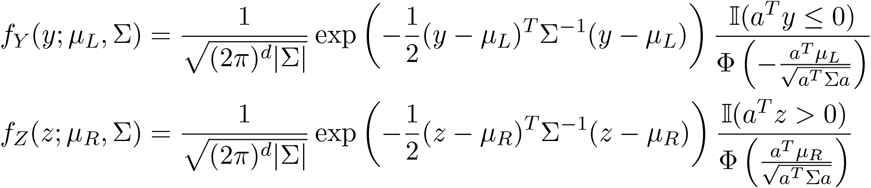

where *μ_L_, μ_R_*, Σ denote the means and covariance matrix of the untruncated versions of the distributions. We assume that all samples are drawn independently, and Σ is diagonal: 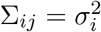 if *i* = *j* else Σ_*ij*_ = 0. The joint distribution of our *n* samples of *Y* with our *m* samples of *Z* can be expressed in exponential family form as

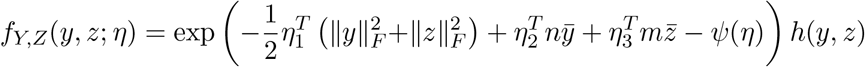

where *ψ* is the cumulant generating function, *h* is some carrying density, and ||·||_*F*_ denotes the Frobenius norm. The natural parameters *η*_1_, *η*_2_, *η*_3_ are equal to

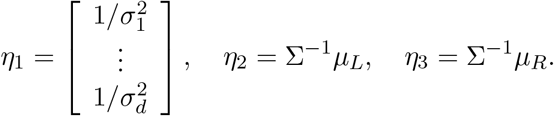

To test differential expression of gene *g*, we can test if *η*_2_*g*__ = *η*_3_*g*__, which is equivalent to testing 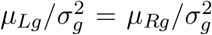 or *μ_L_g__* = *μ_R_g__*. In similar spirit to the 1-dimensional case, we perform a slight reparameterization, letting *θ_g_* = (*η*_3_*g*__ − *η*_2_*g*__)/2 and *μ_g_* = (*η*_2_*g*__ + *η*_3_*g*__)/2:

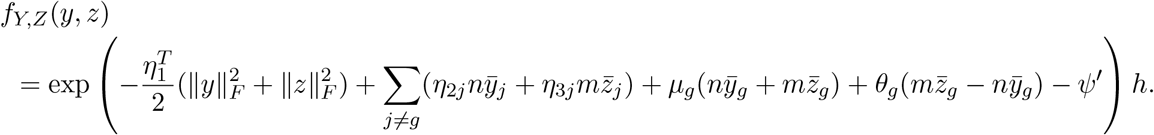

We again design tests for *θ_g_* using its sufficient statistic, 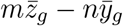:

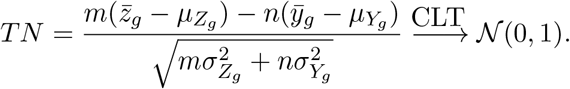

During the testing procedure, we want to evaluate if *θ_g_* =0 (i.e. if gene *g* has significantly different mean expression between the two populations). With *θ_g_* = 0 as our null hypothesis, we compute the corresponding parameters 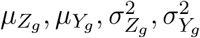 under the null, allowing us to evaluate the probability of seeing a *TN* statistic at least as extreme as the one observed for the actual data.

Like in the 1-dimensional case, we use maximum likelihood to estimate *η*_1_, *η*_2_, and *η*_3_, leveraging the fact that the likelihood function is concave because the joint distribution is an exponential family. After estimating the natural parameters, we can easily recover Σ, *μ_L_*, and *μ_R_*. To obtain estimates for 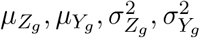 under the null, we first set *μ_L_g__* = *μ_R_g__* = (*μ_L_g__* + *μ_R_g__*)/2. We then use numerical integration to obtain the first and second moments of gene *g*’s marginal distributions:

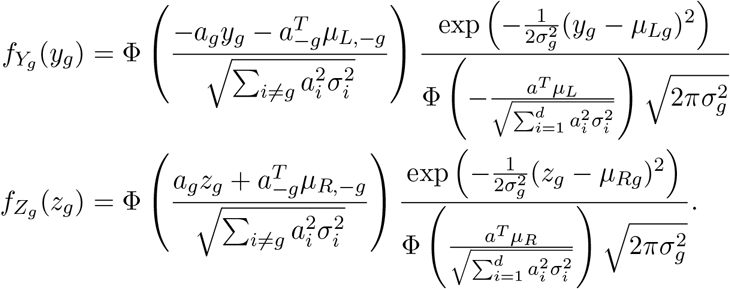

The TN test procedure is summarized in Algorithm 2 and Fig. 1. More details regarding the above derivations are given in the Method S1. Just like in the 1D case, this test is approximate because the tails of our test statistic’s null distribution should be bigger in order to capture the estimation and approximation uncertainty; however, we can obtain significant selection bias correction (for both real and synthetic datasets).

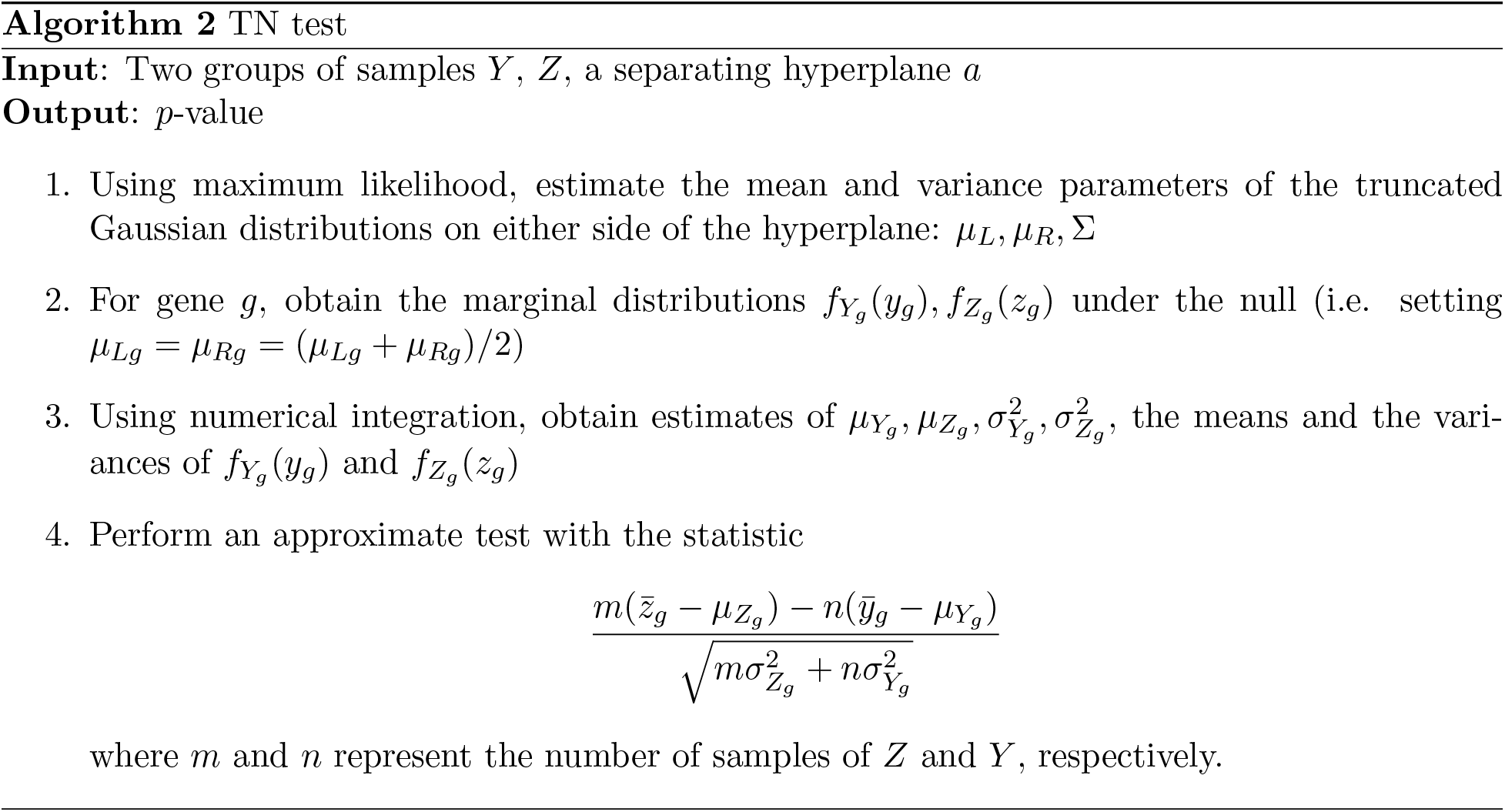

#### TN test for post-clustering *p*-value correcting

We describe a full framework (Fig. 1) for clustering the dataset *X* and obtaining corrected *p*-values via the TN test. Using a data-splitting approach, we run some clustering algorithm on one portion of the data, *X*_1_, to generate 2 clusters. For differential expression analysis, we estimate the separating hyperplane *a* using a linear binary classifier such as the support vector machine (SVM). This hyperplane is used to assign labels to the remaining samples in *X*_2_, yielding *Y* and *Z*. Finally, we can run a TN test using *Y, Z*, and *a*. This approach is summarized in Algorithm 3. Note that in the case of *k* > 2 clusters, we can assign all points in *X*_2_ using our collection of 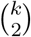 hyperplanes.

##### Algorithm 3 Clustering and TN test framework

~~~
**Input**: Samples *X*
**Output**: *p*-value
    1. Split *X* into two partitions *X*_1_, *X*_2_
    2. Run your favorite clustering algorithm on *X*_1_ to generate labels, choosing two clusters for downstream differential expression analysis
    3. Use *X*_1_ and the labels to determine *a*, the separating hyperplane (e.g. using an SVM)
    4. Divide *X*_2_ into *Y, Z* using the obtained hyperplane
    5. Run TN test using *Y, Z, a*
~~~

## Discussion

The post-selection inference problem arose only recently in the age of big data due to a new paradigm of choosing a model after seeing the data. The problem can be described as a two-step process: 1) *selection* of the model to fit the data based on the data, and 2) *fitting* the selected model. The quality of the fitting is assessed based on the *p*-values associated with parameter estimates, but if the null model does not account for the selection event, then the *p*-values are spurious. This problem was first analyzed in 2013 by statisticians in settings such as selection for linear models under squared loss [18, 19]. Practitioners often select a subset of “relevant” features before fitting the linear model. In other words, the practitioner chooses the best model out of 2^*d*^ possible choices (*d* being the number of features), and the quality of fit is hence biased. One needs to account for the selection in order to correct for this bias [18]. Similarly, a single-cell RNA-seq dataset of *n* cells can be divided into 2 clusters in 2^*n*^ ways, biasing the features selected for distinguishing between clusters. In this manuscript, we propose a way to account for this bias.

The proposed analysis framework involves two major components that help ameliorate the data-snooping issue: (i) the data-splitting procedure (to correct for selection bias), and (ii) the TN test formulation (to ensure we use the right null). We can evaluate the amount of correction provided by each component by considering the four frameworks in which we can test for differential expression:

1. Cluster and apply a *t*-test on the same dataset (standard framework);
2. Cluster and apply a TN test on the same dataset (this will require us to perform hyperplane estimation on the same dataset as well);
3. Split the dataset in half, assign cluster labels to the second half using the first half, and perform a *t*-test on the second half; and
4. Split the dataset in half, assign cluster labels to the second half using the first half, and perform a TN-test on the second half (proposed framework).

As shown in Fig. 4a, the data splitting seems more consequential on real datasets such as the PBMC dataset. Designing truncated versions of other tests (such as negative binomial and MAST) would improve component (ii) and thus be promising extensions of the proposed framework.

**Figure 4:**
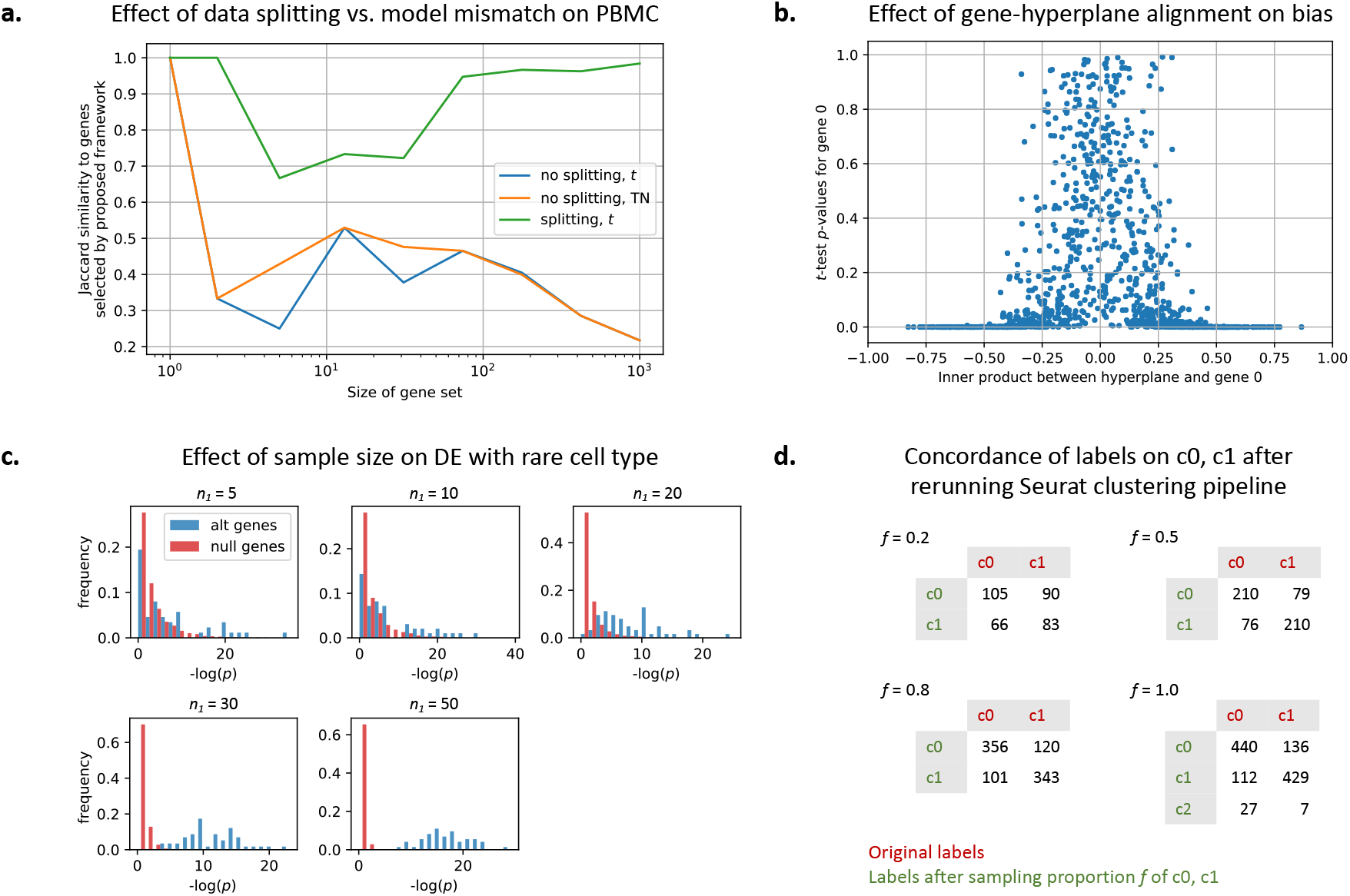
Further experimental exploration of framework. **a.** Comparing the effect of data splitting versus the effect of using the TN test rather than the *t*-test for the PBMC dataset. **b.** A simulation showing the amount of selection bias for a gene with respect to how much the gene aligns with the (given) separating hyperplane. **c.** Detecting differentially expressed genes in a simulated rare cell population of size *n*_1_. The other population has 100 samples. For simulation details, see STAR Methods. **d.** State-of-the-art single-cell clustering pipelines such as Seurat can generate different clustering results on the same cells. Reclustering done for clusters 0 and 1 shown in Figure 2a.

Additionally, we observe that the amount of selection bias a gene suffers is directly related to how much the gene axis intersects the separating boundaries. In this work, the separating boundary is a hyperplane, and therefore the closer to 0 the gene-hyperplane inner product, the less selection bias that gene experiences (Fig. 4b).

We also gauge the sample size needed to detect differentially expressed genes for rare cell populations (Fig. 4c) using synthetic data. We keep one population at a sample size of 100 and vary *n*_1_, the size of the other population. We see that at *n*_1_ < 20, detecting differentially expressed genes is difficult (the distribution of null genes overlap with the distribution of differentially expressed genes). As we increase *n*_1_, the power increases.

In general, although data splitting reduces the number of samples available for clustering, we see that sacrificing a portion of the data can correct for biases introduced by clustering. State-of-the-art single-cell clustering pipelines such as Seurat can generate different clustering results on the same dataset (Fig. 4d). Different clustering results imply different null hypotheses when we reach the differential expression analysis step, which further undermines the validity of the “discovered” differentiating markers. Therefore trading off samples to correct the selection bias may well be worth it.

This work introduced and validated the TN test framework in the single-cell RNA-Seq application, but the framework is equally applicable to other domains where feature sets are large and clustering is done before feature selection. Because science has entered a big-data era where obtaining large datasets is becoming increasingly cheaper, researchers across domains have fallen into the mindset of forming hypotheses after seeing the data [12]. We believe that the TN test is a step towards the right direction: correcting data snooping to reduce false discoveries and improve reproducibility.

## Supporting information

Supplementary figures and additional derivations

## Acknowledgements

We thank Jonathan Taylor, Martin Zhang, and Vasilis Ntranos of Stanford University and Aaron Lun of the Cancer Research UK Cambridge Institute for helpful discussions about selective inference and applications of the method. GMK and JMZ are supported by the Center for Science of Information, an NSF Science and Technology Center, under grant agreement CCF-0939370. JMZ and DNT are supported in part by the National Human Genome Research Institute of the National Institutes of Health under award number R01HG008164.

## Author Contributions

JMZ observed the *p*-value problem for single-cell RNA-Seq computational pipelines. GMK noted the selection bias and conceived the idea of leveraging the post-selection inference framework. GMK and JMZ formulated a solution and designed the experiments. JMZ performed the derivations, implemented the Python module, performed the experiments, and wrote the manuscript. JMZ, GMK, and DNT interpreted results. DNT supervised the project. All authors read and approved the final manuscript.

## STAR⋆ Methods

### Method Details

Please see the Results section for details on the TN test framework and formulation. Additional derivations can be found in the Method S1.

#### Method validation via simulation

To validate the method, we use synthetic datasets where the ground truth is fixed and known. We first show that the TN tests generates valid *p*-values. For the experiments discussed in Supplementary Fig. 1, we sample data from normal distributions with identity covariance prior to clustering, resulting in data sampled from truncated normal distributions post-clustering. To estimate the separating hyperplane *a*, we fit an SVM to 10% of the dataset (50% for Supplementary Fig. 1c), and we work with the remaining portion of the dataset after relabeling it based on our estimate of *a*. Supplementary Fig. 1a shows results for the 2-gene case where no differential expression should be observed (i.e. the untruncated means are identical). Note that for this example, gene 1 needs a larger correction factor than gene 2 because the separating hyperplane is less aligned with the gene 1 axis. We see that when both the variance and separating hyperplane *a* are known, the TN test completely corrects for the selection event. As we introduce more uncertainty (i.e. if we need to estimate variance or *a* or both), the correction factor shrinks; however, the gap is still significantly better than for the *t*-test case. Supplementary Fig. 1b repeats the experiment for the case where gene 1 is differentially expressed. The TN test again corrects for the selection bias in gene 2, but we still obtain significant *p*-values for gene 1 though not nearly as extreme as for the *t*-test case.

Supplementary Fig. 1c shows that as we increase *d*, the number of genes, the minimum TN test *p*-value across all *d* genes follows the family-wise error rate (FWER) curve. Since FWER represents the probability of making at least 1 false discovery and naturally increases with *d*, this highlights the validity of the TN test. In comparison, the *t*-test returns extreme *p*-values especially for lower values of *d*. As *d* increases, however, the selection bias incurred by our simple clustering approach disappears. While the TN test provides less gain in higher (≥ 200) dimensions, we note that for real datasets, cluster identities are often driven by an effectively small amount of genes, which is why several single-cell pipelines perform dimensionality reduction before clustering. Supplementary Fig. 1b and 1d show that when certain genes are differentially expressed, the TN test is still able to find them. For Supplementary Fig. 1d, the data consists of 200 samples, 100 from 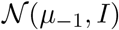 and 100 from 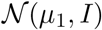 where *μ_i_* ∈ ℝ^500^ is 0 for 490 entries and *i* ∈ {−1,1} for the last 10 entries. This experimental setup was also used for Fig. 4c.

#### Single-cell dataset computational details

For all experiments discussed in Fig. 2 and Fig. 3, we randomly split the set of samples in half into datasets 1 and 2. For Fig. 2a, we recluster dataset 1 with Seurat using clustering parameters that would result in 2 clusters. We use SVM to obtain a hyperplane that perfectly separates the two clusters, and we use this hyperplane to assign labels to samples in dataset 2. When comparing the TN test results to those obtained using other approaches, we run the entire Seurat pipeline (including differential expression analysis) on dataset 1. For the mouse brain cell and mESC datasets analyzed in Fig. 3, we assume that the generated labels are ground truth, and therefore we do not perform the reclustering part of the analysis framework shown in Fig. 1. For Fig. 3a, we only report correction factors for cases where SVM fit the data well, meaning that the new labels generated for dataset 2 have at least a 80% match with the original labels. We note that this does not contradict the linear separability assumption discussed in the main text. The sizes of the interneuron subclusters range from 10 to 26, and therefore the SVM was occasionally fit on as few as 5 samples, resulting in an inability to generalize. Additionally, we only report correction factors greater than 0.

#### Computational Cost

For the PBMC experiment (Fig. 2), comparing two clusters took approximately 3.5 hours on 1 core, the bulk of which was spent performing numerical integration on the marginal distributions for each of over 12000 genes. This process can be sped up by parallelizing the numerical integration on multiple cores or by only processing the genes with the smallest *t*-test *p*-values after estimating the hyperplane using all genes. We provide both options in our software package. In general, estimating the parameters of the truncated normal distribution scales linearly with the dimensionality of the data because we assume that the covariance matrix is diagonal. The estimation problem is straightforward to solve because the joint distribution can be expressed in exponential family form, and therefore gradient methods will converge to a global optimum.

#### Joint distributions of samples (1-dimensional, unit variance case)

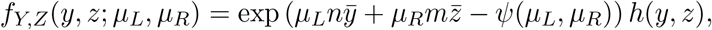

where *ȳ* and 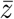 represent the sample means of *Y* and *Z*, respectively. *ψ* is the cumulant generating function, and *h* is the carrying density:

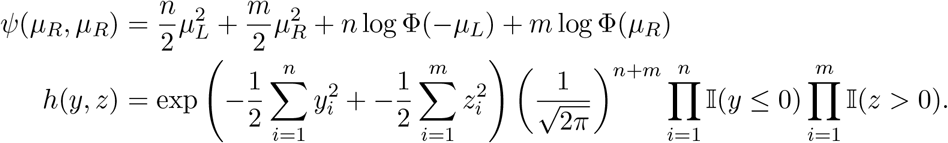

After the reparametrization where we let *θ* = (*μ_R_ − μ_L_*)/2 and *μ* = (*μ_R_* + *μ_L_*)/2, we obtain:

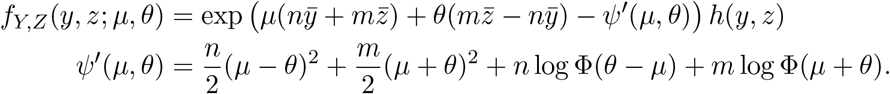

### Quantification and Statistical Analysis

Aside from the statistical methods developed in this manuscript, we used Seurat [6]. We followed the PBMC tutorial provided by the authors at https://satijalab.org/seurat/v3.0/pbmc3k_tutorial.html, deviating only to test other provided differential expression methods. We briefly describe the Seurat differential expression methods presented in Figure 2:

1. Welch’s *t*: a commonly-used variant on the Student’s *t*-test where the variances of the two populations are not assumed to be equal
2. wilcox: the Wilcoxon rank sum test (Seurat’s default test)
3. tobit: Tobit-test for differential gene expression as described by [9]
4. MAST: GLM-framework that treates cellular detection rate as a covariate [14]
5. poisson: likelihood ratio test assuming an underlying Poisson distribution
6. negbinom: likelihood ratio test assuming an underlying negative binomial distribution
7. bimod: likelihood-ratio test for single cell gene expression as described by [13]

In Fig. 2b and Fig. 4a, we quantify overlap using Jaccard similarity, the ratio of the intersection to the union of two sets (a value bounded between 0 and 1).

## Data and Software Availability

Both the software package and the code used to generate the results presented in this paper are available online at https://github.com/jessemzhang/tn_test. The software package can also be installed via PyPI: https://pypi.org/project/truncated-normal/. All single-cell RNA-Seq datasets analyzed were generated from previous studies and can be obtained from public repositories.

## References

[1] Macosko, E. Z. et al. Highly parallel genome-wide expression profiling of individual cells using nanoliter droplets. Cell 161, 1202–1214 (2015).

[2] Zheng, G. X. et al. Massively parallel digital transcriptional profiling of single cells. Nature communications 8, 14049 (2017).

[3] Ntranos, V., Kamath, G. M., Zhang, J. M., Pachter, L. & David, N. T. Fast and accurate single-cell rna-seq analysis by clustering of transcript-compatibility counts. Genome biology 17, 112 (2016).

[4] Buenrostro, J. D. et al. Single-cell chromatin accessibility reveals principles of regulatory variation. Nature 523, 486 (2015).

[5] Habib, N. et al. Massively parallel single-nucleus rna-seq with dronc-seq. Nature methods 14, 955 (2017).

[6] Butler, A., Hoffman, P., Smibert, P., Papalexi, E. & Satija, R. Integrating single-cell transcriptomic data across different conditions, technologies, and species. Nature biotechnology 36, 411 (2018).

[7] Qiu, X. et al. Single-cell mrna quantification and differential analysis with census. Nature methods 14, 309 (2017).

[8] McCarthy, D. J., Campbell, K. R., Lun, A. T. & Wills, Q. F. Scater: pre-processing, quality control, normalization and visualization of single-cell rna-seq data in r. Bioinformatics 33, 1179–1186 (2017).

[9] Trapnell, C. et al. The dynamics and regulators of cell fate decisions are revealed by pseudotemporal ordering of single cells. Nature biotechnology 32, 381 (2014).

[10] Wolf, F. A., Angerer, P. & Theis, F. J. Scanpy: large-scale single-cell gene expression data analysis. Genome biology 19, 15 (2018).

[11] Love, M. I., Huber, W. & Anders, S. Moderated estimation of fold change and dispersion for rna-seq data with deseq2. Genome biology 15, 550 (2014).

[12] Ioannidis, J. P. Why most published research findings are false. PLoS medicine 2, e124 (2005).

[13] McDavid, A. et al. Data exploration, quality control and testing in single-cell qpcr-based gene expression experiments. Bioinformatics 29, 461–467 (2012).

[14] Finak, G. et al. Mast: a flexible statistical framework for assessing transcriptional changes and characterizing heterogeneity in single-cell rna sequencing data. Genome biology 16, 278 (2015).

[15] Kharchenko, P. V., Silberstein, L. & Scadden, D. T. Bayesian approach to single-cell differential expression analysis. Nature methods 11, 740 (2014).

[16] Zhang, J. M., Fan, J., Fan, H. C., Rosenfeld, D. & David, N. T. An interpretable framework for clustering single-cell rna-seq datasets. BMC bioinformatics 19, 93 (2018).

[17] Student. The probable error of a mean. Biometrika 1–25 (1908).

[18] Berk, R. et al. Valid post-selection inference. The Annals of Statistics 41, 802–837 (2013).

[19] Fithian, W., Sun, D. & Taylor, J. Optimal inference after model selection. arXiv preprint arXiv:1410.2597 (2014).

[20] Zeisel, A. et al. Cell types in the mouse cortex and hippocampus revealed by single-cell rna-seq. Science 347, 1138–1142 (2015).

[21] Xu, C. & Su, Z. Identification of cell types from single-cell transcriptomes using a novel clustering method. Bioinformatics 31, 1974–1980 (2015).

[22] Levine, J. H. et al. Data-driven phenotypic dissection of aml reveals progenitor-like cells that correlate with prognosis. Cell 162, 184–197 (2015).

[23] Blondel, V. D., Guillaume, J.-L., Lambiotte, R. & Lefebvre, E. Fast unfolding of communities in large networks. Journal of statistical mechanics: theory and experiment 2008, P10008 (2008).

[24] Brandt, D. Y. et al. Mapping bias overestimates reference allele frequencies at the hla genes in the 1000 genomes project phase i data. G3: Genes, Genomes, Genetics 5, 931–941 (2015).

[25] D’Acquisto, F. et al. Annexin-1 modulates t-cell activation and differentiation. Blood 109, 1095–1102 (2007).

[26] Stelzer, G. et al. The genecards suite: from gene data mining to disease genome sequence analyses. Current protocols in bioinformatics 54, 1–30 (2016).

[27] Kolodziejczyk, A. A. et al. Single cell rna-sequencing of pluripotent states unlocks modular transcriptional variation. Cell stem cell 17, 471–485 (2015).

[28] Lehmann, E. L. & Romano, J. P. Testing statistical hypotheses (Springer Science & Business Media, 2006).

[29] Biase, F. H., Cao, X. & Zhong, S. Cell fate inclination within 2-cell and 4-cell mouse embryos revealed by single-cell rna sequencing. Genome research 24, 1787–1796 (2014).

[30] Birey, F. et al. Assembly of functionally integrated human forebrain spheroids. Nature 545, 54–59 (2017). URL http://dx.doi.org/10.1038/nature22330.

[31] Buettner, F. et al. Computational analysis of cell-to-cell heterogeneity in single-cell rna-sequencing data reveals hidden subpopulations of cells. Nat Biotech 33, 155–160 (2015). URL http://dx.doi.org/10.1038/nbt.3102.

[32] Deng, Q., Ramsköld, D., Reinius, B. & Sandberg, R. Single-cell rna-seq reveals dynamic, random monoallelic gene expression in mammalian cells. Science 343, 193–196 (2014). URL http://science.sciencemag.org/content/343/6167/193. http://science.sciencemag.org/content/343/6167/193.full.pdf.

[33] Joost, S. et al. Single-cell transcriptomics reveals that differentiation and spatial signatures shape epidermal and hair follicle heterogeneity. Cell systems 3, 221–237 (2016).

[34] Patel, A. P. et al. Single-cell rna-seq highlights intratumoral heterogeneity in primary glioblastoma. Science 344, 1396–1401 (2014). URL http://science.sciencemag.org/content/344/6190/1396. http://science.sciencemag.org/content/344/6190/1396.full.pdf.

[35] Pollen, A. A. et al. Low-coverage single-cell mrna sequencing reveals cellular heterogeneity and activated signaling pathways in developing cerebral cortex. Nat Biotech 32, 1053–1058 (2014). URL http://dx.doi.org/10.1038/nbt.2967.

[36] Ting, D. T. et al. Single-cell {RNA} sequencing identifies extracellular matrix gene expression by pancreatic circulating tumor cells. Cell Reports 8, 1905–1918 (2014). URL http://www.sciencedirect.com/science/article/pii/S2211124714007050.

[37] Treutlein, B. et al. Reconstructing lineage hierarchies of the distal lung epithelium using singlecell rna-seq. Nature 509, 371–375 (2014). URL http://dx.doi.org/10.1038/nature13173.

[38] Usoskin, D. et al. Unbiased classification of sensory neuron types by large-scale single-cell rna sequencing. Nat Neurosci 18, 145–153 (2015). URL http://dx.doi.org/10.1038/nn.3881.

[39] Yan, L. et al. Single-cell rna-seq profiling of human preimplantation embryos and embryonic stem cells. Nat Struct Mol Biol 20, 1131–1139 (2013). URL http://dx.doi.org/10.1038/nsmb.2660.

