## Supplementary figures and additional derivations for "Valid post-clustering differential analysis for single-cell RNA-Seq"

#### Supplementary Material

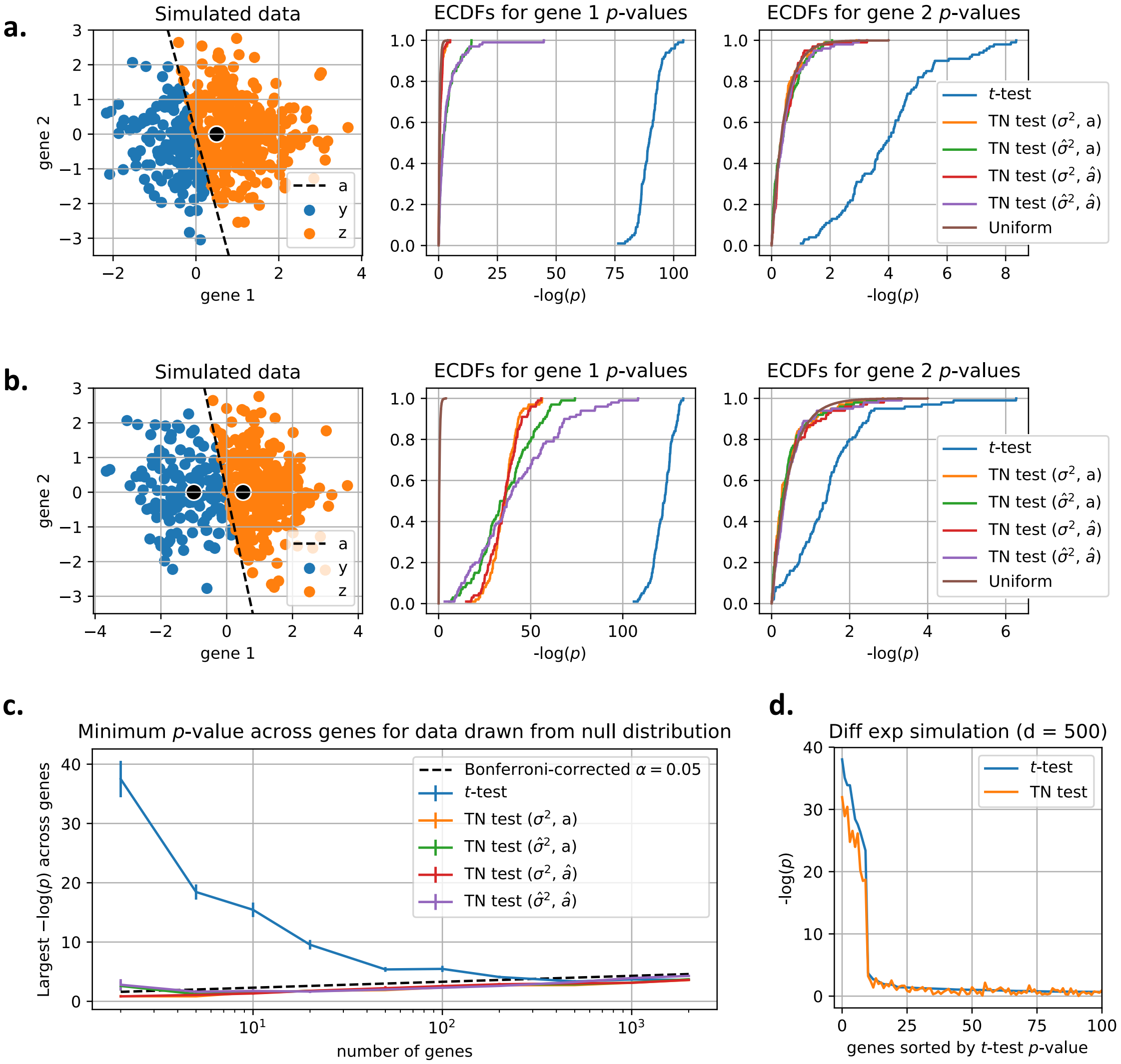


Figure 1: **Results on simulated data drawn from truncated normal distributions, Related to Results. a.** 500 samples are drawn from the same distribution, and genes 1 and 2 are drawn from $\mathcal{N}(0.5, 1)$ and $\mathcal{N}(0, 1)$, respectively. The clustering step splits the dataset into groups of 156 and 344 samples, and $a$ exactly captures the clustering rule. We see that although neither gene is differentially expressed in the underlying distribution, the $t$-test consistently returns small $p$-values across 100 simulation runs. We present four versions of the TN test, all of which significantly correct for the clustering step. $\hat{\sigma}^2$ indicates that the variance was unknown and therefore estimated from the data. $\hat{a}$ indicates that the hyperplane was estimated from a held-out 10% of the samples using an SVM. **b.** The experiment from **a** is repeated except gene 1 is drawn from a $\mathcal{N}(-1, 0)$ distribution instead for one of the clusters. The number of samples in each group and the separating hyperplane remain the same. **c.** We explore how the minimum $p$-value across genes changes with $d$, the number of genes. For a particular number of genes, 200 samples are drawn from a $\mathcal{N}(0, I)$ distribution, and $a$ is chosen randomly. This simulation is repeated 10 times for each value of $d$. $\alpha$ indicates the chosen level of significance. **d.** For $d = 500$, we run a 200-sample simulation experiment (100 in each cluster) where 10 genes are differentially expressed. 10 values of $\mu_L$ were set to -1, and the corresponding entries in $\mu_R$ were set to 1. All other entires of $\mu_L, \mu_R$ were set to 0, and $\sigma^2 = 1$.

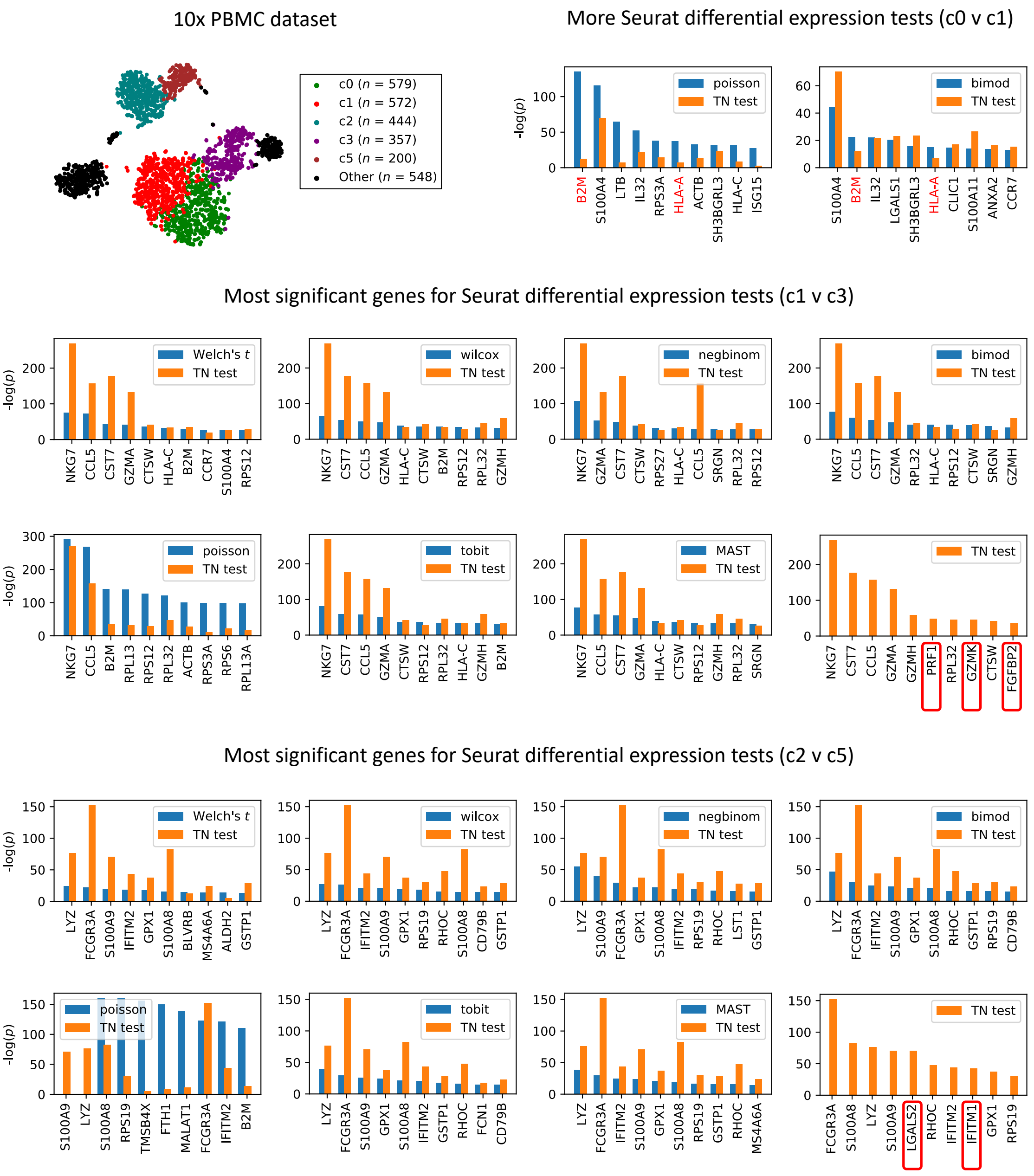


Figure 2: **Comparison of differential expression tests on PBMC dataset continued, Related to Figure 2a.** The comparison performed in Figure 2a is repeated for other Seurat differential expression methods and for clusters 1 v 3 and 2 v 5. A missing bar indicates a $p$ value of 0 due to numerical precision limitations. TN test genes boxed in red were missed by the other tests.

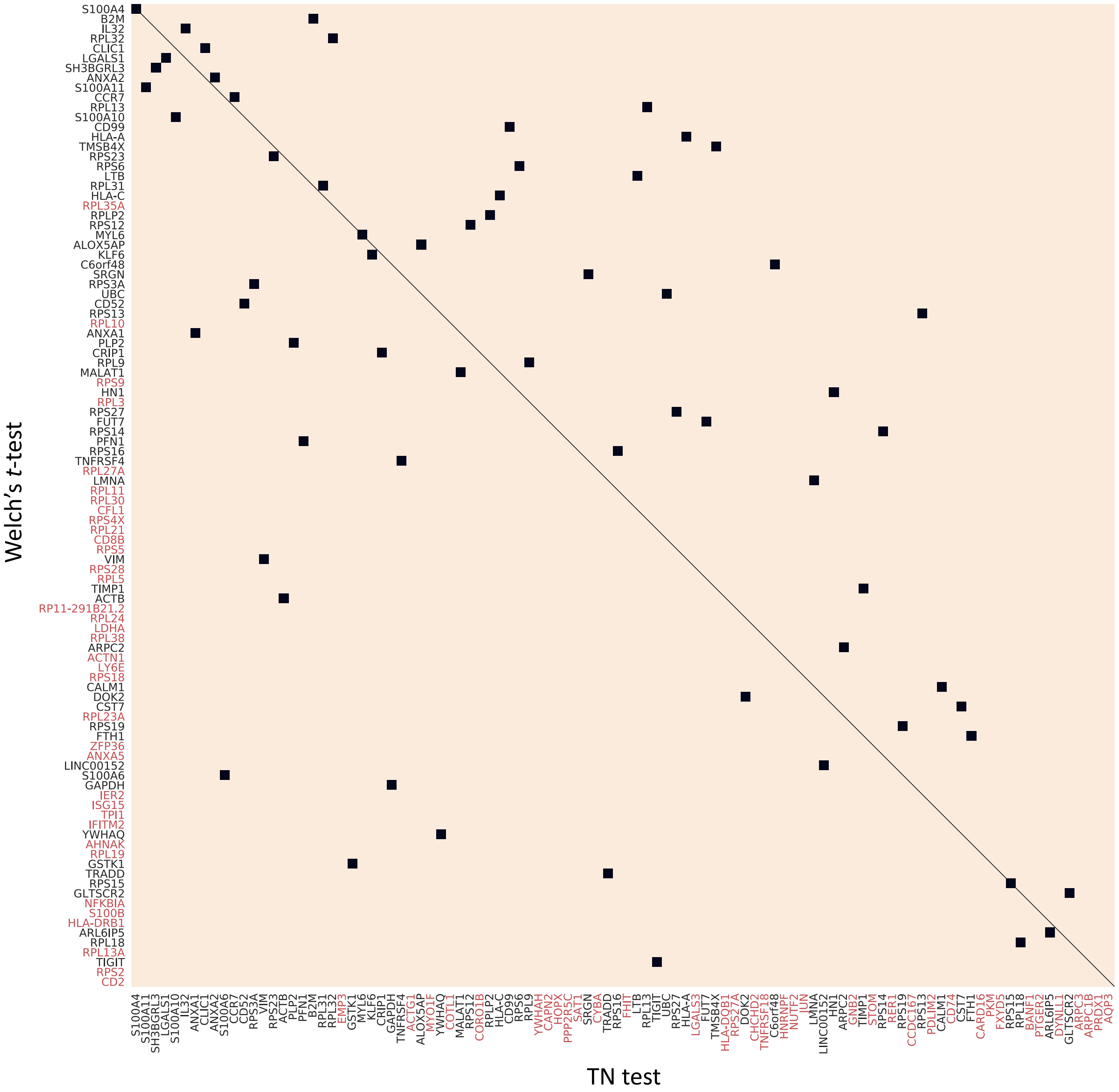


Figure 3: **Gene set comparison using permutation matrix, Related to Figure 2a.** The top 100 genes selected by a Welch's $t$-test compared to the top 100 genes selected by the TN test when performing differential expression analysis on clusters 0 and 1 in Fig. 2a. Genes written in red text were not selected by the other test. 64 genes appear in both gene sets.

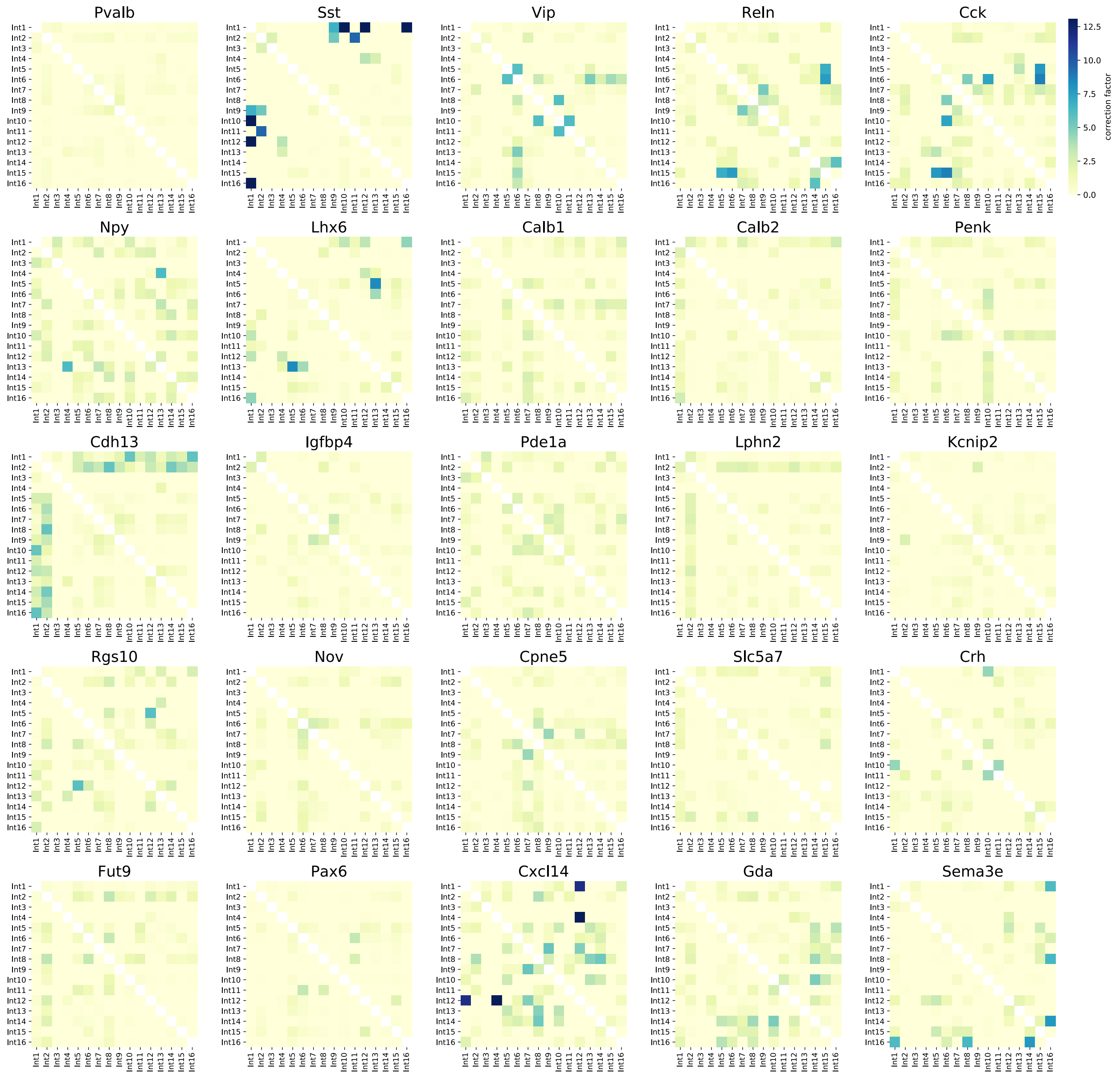


Figure 4: **TN test correction for mouse brain cell dataset interneuron genes, Related to Figure 3a.** The 16 interneuron subclasses reported for the mouse brain cell dataset [20] are re-compared pairwise using each of the 26 genes discussed by the authors. For each gene and pair of subclasses, the correction factor represents the -log of the ratio of the $t$-test $p$-value to the TN test $p$-value. We only consider comparisons where the hyperplane fit the data relatively well (58.3% of comparisons).

### Method S1

#### Joint distributions of samples (multidimensional, diagonal covariance case)

We assume we have $n$ samples of $Y_i$ and $m$ samples of $Z_i$, and we assume that the two populations share the same covariance matrix $\Sigma$. Additionally, we assume we have $d$ genes (i.e. $\Sigma \in \mathbb{R}^{d\times d}$). We again let $a$ represent our separating hyperplane, and we include $b$ as our intercept term. We can compute $G_a(\mu, \Sigma)$, the normalization factor for our multivariate truncated normal distribution, as

$$
\begin{aligned}
G_a(\mu,\Sigma) &= P(a^T X + b > 0), && X \sim \mathcal{N}(\mu,\Sigma)\\
&= P(\sqrt{a^T\Sigma a} Z + a^T\mu + b > 0), && Z \sim \mathcal{N}(0,1)\\
&= P\left(z > \frac{-a^T\mu - b}{\sqrt{a^T\Sigma a}}\right)\\
&= \Phi\left(\frac{a^T\mu + b}{\sqrt{a^T\Sigma a}}\right),
\end{aligned}
$$

overloading our notation a bit to let $Z$ represent a standard normal random variable. Our truncated distributions are

$$
\begin{aligned}
f_Y(y;\mu_L,\Sigma) &= \frac{1}{\sqrt{(2\pi)^d|\Sigma|}} \exp\left(-\frac{1}{2}(y-\mu_L)^T\Sigma^{-1}(y-\mu_L)\right)\frac{\mathbb{I}(a^Ty + b \le 0)}{\Phi\left(\frac{-a^T\mu_L - b}{\sqrt{a^T\Sigma a}}\right)}\\
f_Z(z;\mu_R,\Sigma) &= \frac{1}{\sqrt{(2\pi)^d|\Sigma|}} \exp\left(-\frac{1}{2}(z-\mu_R)^T\Sigma^{-1}(z-\mu_R)\right)\frac{\mathbb{I}(a^Tz + b > 0)}{\Phi\left(\frac{a^T\mu_R + b}{\sqrt{a^T\Sigma a}}\right)},
\end{aligned}
$$

resulting in the joint distribution

$$
f_{Y,Z}(y,z;\eta) = \exp\left(-\frac{1}{2}\sum_{i=1}^n y_i^T\Sigma^{-1}y_i - \frac{1}{2}\sum_{i=1}^m z_i^T\Sigma^{-1}z_i + \mu_L^T\Sigma^{-1}\sum_{i=1}^n y_i + \mu_R^T\Sigma^{-1}\sum_{i=1}^m z_i - \psi(\eta)\right)h(y,z)
$$

$$
h(y,z) = (2\pi)^{\frac{d}{2}(n+m)}\prod_{i=1}^n \mathbb{I}(a^Ty_i + b \le 0)\prod_{i=1}^m \mathbb{I}(a^Tz_i + b > 0)
$$

$$
\psi(\eta) = \frac{n}{2}\mu_L^T\Sigma^{-1}\mu_L + \frac{m}{2}\mu_R^T\Sigma^{-1}\mu_R + n\log\Phi\left(\frac{-a^T\mu_L - b}{\sqrt{a^T\Sigma a}}\right) + m\log\Phi\left(\frac{a^T\mu_R + b}{\sqrt{a^T\Sigma a}}\right) + \frac{n+m}{2}\log|\Sigma|.
$$

#### Maximum likelihood estimation of parameters via gradient ascent

As discussed in the main text, we assume $\Sigma$ is diagonal: $\Sigma_{ij} = \sigma_i^2$ if $i = j$ else $\Sigma_{ij} = 0$. We reparametrize the joint distribution in exponential family form as

$$
f_{Y,Z}(y,z;\eta) = \exp\left(-\frac{1}{2}\eta_1^T\left(\sum_{i=1}^n y_i \circ y_i + \sum_{i=1}^m z_i \circ z_i\right) + \eta_2^T\sum_{i=1}^n y_i + \eta_3^T\sum_{i=1}^m z_i - \psi(\eta)\right)h(y,z)
$$

where $\circ$ represents element-wise multiplication, and the natural parameters $\eta_1, \eta_2, \eta_3$ are equal to

$$\eta_1 = \begin{bmatrix} 1/\sigma_1^2 \\ \vdots \\ 1/\sigma_d^2 \end{bmatrix}, \quad \tilde{\eta}_1 = \begin{bmatrix} \sigma_1^2 \\ \vdots \\ \sigma_d^2 \end{bmatrix}, \quad \eta_2 = \Sigma^{-1}\mu_L, \quad \eta_3 = \Sigma^{-1}\mu_R,$$

$h$ remains the same as above, but $\psi$ is now

$$\begin{aligned}
\psi(\eta) &= \frac{n}{2}\eta_2^T(\tilde{\eta}_1 \circ \eta_2) + \frac{m}{2}\eta_3^T(\tilde{\eta}_1 \circ \eta_3) + n \log \Phi(c_Y) + m \log \Phi(c_Z) + \frac{n+m}{2} \log|\tilde{\eta}_1| \\
c_Y &= \frac{-a^T(\tilde{\eta}_1 \circ \eta_2) - b}{\sqrt{a^T(\tilde{\eta}_1 \circ a)}} \\
c_Z &= \frac{a^T(\tilde{\eta}_1 \circ \eta_3) + b}{\sqrt{a^T(\tilde{\eta}_1 \circ a)}}.
\end{aligned}$$

We use maximum likelihood (ML) to estimate $\eta_1, \eta_2$, and $\eta_3$, leveraging the fact that the likelihood function is concave because the joint distribution is an exponential family. Because we cannot express our ML estimators in closed form, we instead use gradient ascent. The gradient update equations can be derived to be

$$\begin{aligned}
\frac{\partial \ell}{\partial \eta_2} &= \sum_{i=1}^{n} y_i - n\tilde{\eta}_1 \circ \eta_2 + n\frac{\phi(c_Y)}{\Phi(c_Y)} \frac{\tilde{\eta}_1 \circ a}{\sqrt{a^T(\tilde{\eta}_1 \circ a)}} \\
\frac{\partial \ell}{\partial \eta_3} &= \sum_{i=1}^{m} z_i - m\tilde{\eta}_1 \circ \eta_3 - m\frac{\phi(c_Z)}{\Phi(c_Z)} \frac{\tilde{\eta}_1 \circ a}{\sqrt{a^T(\tilde{\eta}_1 \circ a)}} \\
\frac{\partial \ell}{\partial \eta_1} &= -\frac{1}{2}\sum_{i=1}^{n} y_i \circ y_i - \frac{1}{2}\sum_{i=1}^{m} z_i \circ z_i + \tilde{\eta}_1 \circ \tilde{\eta}_1 \circ \left[\frac{n\eta_2 \circ \eta_2 + m\eta_3 \circ \eta_3 + (n+m)\eta_1}{2}\right] \\
&\quad - \tilde{\eta}_1 \circ \tilde{\eta}_1 \circ \left[\frac{n}{2}\frac{\phi(c_Y)}{\Phi(c_Y)} \left( \frac{2a \circ \eta_2}{\sqrt{a^T(\tilde{\eta}_1 \circ a)}} - \frac{a^T(\tilde{\eta}_1 \circ \eta_2) + b}{\left(\sqrt{a^T(\tilde{\eta}_1 \circ a)}\right)^3} a \circ a \right)\right] \\
&\quad - \tilde{\eta}_1 \circ \tilde{\eta}_1 \circ \left[\frac{m}{2}\frac{\phi(c_Z)}{\Phi(c_Z)} \left( \frac{a^T(\tilde{\eta}_1 \circ \eta_3) + b}{\left(\sqrt{a^T(\tilde{\eta}_1 \circ a)}\right)^3} a \circ a - \frac{2a \circ \eta_3}{\sqrt{a^T(\tilde{\eta}_1 \circ a)}} \right)\right]
\end{aligned}$$

where quantities inside brackets are $d$-dimensional vectors. After obtaining the estimates for $\eta_1, \eta_2, \eta_3$, we can obtain estimates for the original parameters:

$$\begin{aligned}
\hat{\Sigma} &= \mathrm{diag}(\hat{\tilde{\eta}}_1) \\
\hat{\mu}_L &= \hat{\tilde{\eta}}_1 \circ \hat{\eta}_2 \\
\hat{\mu}_R &= \hat{\tilde{\eta}}_1 \circ \hat{\eta}_3.
\end{aligned}$$

#### Marginal distributions for a particular gene $g$

Without loss of generality, we consider $g = 1$. The following substitution will be useful when computing the marginal distributions of $Y_1$ and $Z_1$:

$$V = \begin{bmatrix} V_{1,1} & V_{1,-1} \\ V_{-1,1} & V_{-1,-1} \end{bmatrix} = \Sigma^{-1}.$$

We also use the $-1$ subscript to indicate "all indices except the first." We start with the distribution of $Y$:

$$\begin{aligned} f_Y(y;\mu_L,\Sigma) &= \frac{1}{\sqrt{(2\pi)^d|\Sigma|}} \exp\left(-\frac{1}{2}(y-\mu_L)^T\Sigma^{-1}(y-\mu_L)\right) \frac{\mathbb{I}(a^Ty+b\leq 0)}{\Phi\left(\frac{-a^T\mu_L-b}{\sqrt{a^T\Sigma a}}\right)} \\ &= \frac{\mathbb{I}(a^Ty+b\leq 0)}{C} \exp\left(-\frac{1}{2}(y_1-\mu_{L1})^2V_{1,1} - \frac{1}{2}(y_{-1}-\mu_{L,-1})^TV_{-1,-1}(y_{-1}-\mu_{L,-1})\right) \\ &\qquad \exp\left(-(y_1-\mu_{L1})V_{1,-1}(y_{-1}-\mu_{L,-1})\right). \end{aligned}$$

We can complete the square here using the identity:

$$\frac{1}{2}x^TAx + b^Tx + c = \frac{1}{2}(x+A^{-1}b)^TA(x+A^{-1}b) + c - \frac{1}{2}b^TA^{-1}b$$

and letting

$$\begin{aligned} x &= y_{-1}-\mu_{L,-1} \\ A &= V_{-1,-1} \\ b &= V_{-1,1}(y_1-\mu_{L1}) \\ c &= \frac{1}{2}(y_1-\mu_{L1})^2V_{1,1}, \end{aligned}$$

resulting in

$$\begin{aligned} f_Y(y;\mu_L,\Sigma) = \frac{\mathbb{I}(a^Ty+b\leq 0)}{C} &\exp\left(-\frac{1}{2}(y_1-\mu_{L1})^2(V_{1,1}-V_{1,-1}V_{-1,-1}^{-1}V_{-1,1})\right) \\ &\exp\left(-\frac{1}{2}[y_{-1}-\mu_{L,-1}+V_{-1,-1}^{-1}V_{-1,1}(y_1-\mu_{L1})]^TV_{-1,-1}[y_{-1}-\mu_{L,-1}+V_{-1,-1}^{-1}V_{-1,1}(y_1-\mu_{L1})]\right). \end{aligned}$$

If we now marginalize out $y_{-1}$, we get

$$f_{Y_1}(y_1) = \frac{1}{C}\exp\left(-\frac{1}{2}(y_1-\mu_{L1})^2(V_{1,1}-V_{1,-1}V_{-1,-1}^{-1}V_{-1,1})\right) P(a^TY+b\leq 0|Y_1=y_1)\sqrt{(2\pi)^{d-1}|V_{-1,-1}^{-1}|}.$$

We use the fact that $Y_{-1}|Y_1 = y_1 \sim \mathcal{N}\left(\mu_{L,-1} - V_{-1,-1}^{-1}V_{-1,1}(y_1-\mu_{L1}), V_{-1,-1}^{-1}\right) = \mathcal{N}(\tilde{\mu},\tilde{\Sigma})$ to show that

$$\begin{aligned} P(a^T Y + b \le 0 | Y_1 = y_1) &= P(a_1 y_1 + a_{-1}^T Y_{-1} + b \le 0) \\ &= P\left(a_1 y_1 + a_{-1}^T \tilde{\mu} + \sqrt{a_{-1}^T \tilde{\Sigma} a_{-1}} X + b \le 0\right), \qquad X \sim \mathcal{N}(0,1) \\ &= P\left(X \le \frac{-a_1 y_1 - a_{-1}^T \tilde{\mu} - b}{\sqrt{a_{-1}^T \tilde{\Sigma} a_{-1}}}\right) \\ &= \Phi\left(\frac{-a_1 y_1 - a_{-1}^T \tilde{\mu} - b}{\sqrt{a_{-1}^T \tilde{\Sigma} a_{-1}}}\right). \end{aligned}$$

Therefore,

$$f_{Y_1}(y_1) = \Phi\left(\frac{-a_1 y_1 - a_{-1}^T(\mu_{L,-1} - V_{-1,-1}^{-1} V_{-1,1}(y_1 - \mu_{L1})) - b}{\sqrt{a_{-1}^T V_{-1,-1}^{-1} a_{-1}}}\right) \frac{\exp\left(-\frac{1}{2}(y_1 - \mu_{L1})^2 (V_{1,1} - V_{1,-1} V_{-1,-1}^{-1} V_{-1,1})\right)}{\Phi\left(\frac{-a^T \mu_L - b}{\sqrt{a^T \Sigma a}}\right) \sqrt{2\pi |\Sigma| |V_{-1,-1}|}}.$$

We can repeat the process for $Z$, which has distribution

$$f_Z(z; \mu_R, \Sigma) = \frac{1}{\sqrt{(2\pi)^d |\Sigma|}} \exp\left(-\frac{1}{2}(y - \mu_R)^T \Sigma^{-1} (z - \mu_R)\right) \frac{\mathbb{I}(a^T z + b > 0)}{\Phi\left(\frac{a^T \mu_R + b}{\sqrt{a^T \Sigma a}}\right)},$$

resulting in

$$f_{Z_1}(z_1) = \frac{1}{C} \exp\left(-\frac{1}{2}(z_1 - \mu_{R1})^2 (V_{1,1} - V_{1,-1} V_{-1,-1}^{-1} V_{-1,1})\right) P(a^T Z + b > 0 | Z_1 = z_1) \sqrt{(2\pi)^{d-1} |V_{-1,-1}^{-1}|}.$$

We evaluate the probability term as

$$\begin{aligned} P(a^T Z + b > 0 | Z_1 = z_1) &= P(a_1 z_1 + a_{-1}^T Z_{-1} + b > 0) \\ &= P\left(a_1 z_1 + a_{-1}^T \tilde{\mu} + \sqrt{a_{-1}^T \tilde{\Sigma} a_{-1}} X + b > 0\right), \qquad X \sim \mathcal{N}(0,1) \\ &= P\left(X > \frac{-a_1 z_1 - a_{-1}^T \tilde{\mu} - b}{\sqrt{a_{-1}^T \tilde{\Sigma} a_{-1}}}\right) \\ &= \Phi\left(\frac{a_1 z_1 + a_{-1}^T \tilde{\mu} + b}{\sqrt{a_{-1}^T \tilde{\Sigma} a_{-1}}}\right), \end{aligned}$$

resulting in the marginal distribution

$$f_{Z_1}(z_1) = \Phi\left(\frac{a_1 z_1 + a_{-1}^T(\mu_{R,-1} - V_{-1,-1}^{-1} V_{-1,1}(z_1 - \mu_{R1})) + b}{\sqrt{a_{-1}^T V_{-1,-1}^{-1} a_{-1}}}\right) \frac{\exp\left(-\frac{1}{2}(z_1 - \mu_{R1})^2 (V_{1,1} - V_{1,-1} V_{-1,-1}^{-1} V_{-1,1})\right)}{\Phi\left(\frac{a^T \mu_R + b}{\sqrt{a^T \Sigma a}}\right) \sqrt{2\pi |\Sigma| |V_{-1,-1}|}}.$$

Because we assume diagonal covariance, the above marginal distributions can be simplified to:

$$f_{Y_1}(y_1) = \Phi\left(\frac{-a_1 y_1 - a_{-1}^T \mu_{L,-1} - b}{\sqrt{\sum_{i=2}^d a_i^2 \sigma_i^2}}\right) \frac{\exp\left(-\frac{1}{2\sigma_1^2}(y_1 - \mu_{L1})^2\right)}{\Phi\left(\frac{-a^T \mu_L - b}{\sqrt{\sum_{i=1}^d a_i^2 \sigma_i^2}}\right)\sqrt{2\pi\sigma_1^2}}$$

$$f_{Z_1}(z_1) = \Phi\left(\frac{a_1 z_1 + a_{-1}^T \mu_{R,-1} + b}{\sqrt{\sum_{i=2}^d a_i^2 \sigma_i^2}}\right) \frac{\exp\left(-\frac{1}{2\sigma_1^2}(z_1 - \mu_{R1})^2\right)}{\Phi\left(\frac{a^T \mu_R + b}{\sqrt{\sum_{i=1}^d a_i^2 \sigma_i^2}}\right)\sqrt{2\pi\sigma_1^2}}.$$
